# Long-lasting contribution of dopamine in the nucleus accumbens core, but not dorsal lateral striatum, to sign-tracking

**DOI:** 10.1101/132977

**Authors:** Kurt M. Fraser, Patricia H. Janak

**Affiliations:** Department of Psychological and Brain Sciences, Krieger School of Arts and Sciences, Johns Hopkins University, Baltimore, MD 21218, USA; The Solomon H. Snyder Department of Neuroscience, Johns Hopkins School of Medicine, Johns Hopkins University, Baltimore, MD 21205, USA

**Keywords:** incentive salience, reward, Pavlovian conditioning, motivation, rat

## Abstract

The attribution of incentive salience to reward-paired cues is dependent on dopamine release in the nucleus accumbens core. These dopamine signals conform to traditional reward-prediction error signals and have been shown to diminish with time. Here we examined if the diminishing dopamine signal in the nucleus accumbens core has functional implications for the expression of sign-tracking, a Pavlovian conditioned response indicative of the attribution of incentive salience to reward-paired cues. Food-restricted male Sprague-Dawley rats were trained in a Pavlovian paradigm in which an insertable lever predicted delivery of food reward in a nearby food cup. After 7 or 14 training sessions, rats received infusions of saline, the dopamine antagonist flupenthixol (100 mM), or the GABA agonists baclofen and muscimol (0.5 mM baclofen/0.05 mM muscimol) into the nucleus accumbens core or the dorsal lateral striatum. Dopamine antagonism within the nucleus accumbens core attenuated sign-tracking, whereas reversible inactivation did not affect sign-tracking but increased non-specific food cup checking behaviors. Neither drug in the dorsal lateral striatum affected sign-tracking behavior. Critically, extended training did not alter these effects. Though extended experience with an incentive stimulus may reduce cue-evoked dopamine in the nucleus accumbens core, this does not alter the function of dopamine in this region to promote Pavlovian cue approach nor result in the recruitment of dorsal lateral striatal systems for this behavior. These data support the notion that dopamine within the mesoaccumbal system, but not the nigrostriatal system, contributes critically to incentive motivational processes independent of the length of training.

**Abbreviations:**

DLS: dorsal lateral striatum
GT: goal-tracker
IN: intermediate responder
NAcC: nucleus accumbens core
ST: sign-tracker

## 1. Introduction

Environmental stimuli associated with rewards come to guide and direct behavior based on their acquired predictive and incentive motivational properties. Dopamine is critical for both learning the relationship between cues and the outcomes they predict (Schultz et al., 1997; Hollerman and Schultz, 1998; Steinberg et al., 2013), as well as the attribution of incentive motivational value to those cues (Berridge and Robinson, 1998; Berridge, 2007, 2012; Flagel et al., 2011b). However, it remains unclear how dopamine coordinates distinct reward-related processes in distributed brain regions, as well as over a diversity of timescales.

Individual differences in Pavlovian conditioned approach behaviors have been exploited as a means to dissociate aspects of reward learning from incentive motivational processes. Following pairings of a discrete, localizable cue with reward some animals, “sign-trackers” (Hearst and Jenkins, 1974), develop a conditioned response primarily directed towards the cue itself, while others, “goal-trackers” (Boakes, 1977), instead approach and interact with the site of reward delivery upon cue presentation. Only for sign-trackers does the cue acquire properties of an “incentive stimulus” (Berridge, 2001; Cardinal et al., 2002), as sign-trackers approach the cue and will avidly work to obtain its presentation (Robinson and Flagel, 2009). Dopamine release in the nucleus accumbens core (NAcC) transfers with learning from reward receipt to cue presentation and eventually decays with increased experience only for sign-trackers and not goal-trackers (Flagel et al., 2011b; Clark et al., 2013). These dynamics in the NAcC of sign-trackers reflect those predicted by reward-prediction error theories of dopamine’s function (Flagel et al., 2011b; Keiflin and Janak, 2015). Accordingly, blockade of dopamine in the NAcC blunts the expression of sign-tracking, but not goal-tracking (Saunders and Robinson, 2012). Thus dopamine acts in the NAcC to facilitate the attribution of incentive salience to reward-associated stimuli and is necessary for the expression of Pavlovian cue approach, but is not necessary for all forms of Pavlovian reward learning.

Incentive stimuli evoke activity in all regions of the striatum, though little is known about the underlying circuitry of incentive salience attribution apart from investigations into the contributions of the NAcC and its inputs (Flagel and Robinson, 2017). Experience-dependent shifts from ventral medial striatum, including NAcC, to dorsal lateral striatum have been linked to compulsive drug-seeking in rodents and humans (Belin and Everitt, 2008; Vollstädt-Klein et al., 2010; Everitt and Robbins, 2013). Resolving the time-dependency of the systems regulating sign-tracking is critical as sign-tracking renders individuals susceptible to cue-induced craving and seeking for food and drug reward (Saunders and Robinson, 2010, 2013; Saunders et al., 2013). To address this, we explored whether the contribution of neural activity or dopamine in NAcC to sign-tracking is altered with extended experience and additionally if activity or dopamine in dorsal lateral striatum is critical for sign-tracking.

## 2. Methods

### 2.1 Subjects

Adult male Sprague-Dawley rats (initial n=90) approximately 60 days of age, weighing 250-300g on arrival, were obtained from ENVIGO (Barrier 208A; Frederick, MD) and were single-housed with enrichment in a temperature and humidity controlled room maintained on a 12-h light/dark cycle (lights on at 07:00 h). Upon arrival rats were left undisturbed for one week to habituate to the housing environment, during which food and water were available *ad libitum*. Prior to conditioning, rats were mildly food-restricted to 95% of their free-feeding body weight to increase the development of sign-tracking (Anderson et al., 2013) and to match the experimental procedures and design of Clark et al., (2013). Rats were fed 18 g of chow at the conclusion of each day of training and weights were monitored throughout the course of the experiment. All experimental procedures followed recommended guidelines published in the Guide for the Care and Use of Laboratory Animals: Eighth Edition, revised in 2011, in addition to being approved by the Animal Care and Use Committee at The Johns Hopkins University.

### 2.2 Surgery

Prior to behavioral training, all subjects were implanted with bilateral cannula targeted to the dorsal lateral striatum (DLS) or nucleus accumbens core (NAcC) with procedures similar to those previously described (Corbit et al., 2012, 2014). Rats were deeply anesthetized with 5% isoflurane gas and placed in a stereotaxic device (Kopf Instruments; Tujunga, CA) and maintained at 1-2% isoflurane for the duration of surgery. The scalp was shaved and subsequently cleaned with 70% ethyl alcohol and Betadine solution and incised to expose the skull. The skull was cleaned and leveled between bregma and lambda within 0.1 mm of accuracy. A 26 gauge guide cannula (Plastics One; Roanoke, VA) was implanted above the DLS in each hemisphere (AP: +1.2 mm, ML ± 3.4 mm, DV −1.0 mm; all coordinates relative to bregma; n=43) or a 22 gauge cannula (Plastics One; Roanoke, VA) was implanted above the NAcC in each hemisphere (AP: +1.8 mm, ML ± 1.55 mm, DV −5.0 mm; n=47). The cannula gauge and coordinates were based off of previous studies (Corbit and Janak, 2007; Corbit et al., 2012, 2014; Saunders and Robinson, 2012). Cannula were anchored to the skull using three to four bone screws and acrylic dental cement. Stainless steel stylets were inserted in each cannula immediately after surgery and remained in place at all times except during infusions to prevent occlusion. After surgery rats received an injection of cefazolin (70 mg/kg; s.c.) to prevent infection and carprofen (50 mg/kg; s.c.) to alleviate pain. Rats were allowed to recover for at least five days before training began.

### 2.3 Behavioral Apparatus

Training and testing took place in 8 Med Associates (St. Albans, VT) operant chambers housed in sound-and light-attenuating cabinets. In the center of one wall of the chamber was a food cup equipped with a pellet dispenser containing 45 mg banana flavored food pellets (BioServ, Flemington, NJ). The food cup contained a photobeam that recorded beam breaks as food cup entries. On either the left or right side of the food cup was a retractable lever counterbalanced across chambers. When extended, the lever required a 10 g force to record a deflection. A white houselight was positioned on the top of the wall opposite the food cup and lever that provided illumination during each session. Computers with MED-PC software controlled the equipment and recorded lever deflections, latency to first lever contact during a trial, food cup entries, and latency to first food cup entry during a trial.

### 2.4 Pavlovian Conditioning Procedures

The conditioning procedures employed were similar to those that have been described elsewhere (Haight et al., 2015; Fraser et al., 2016) and are diagrammed in Figure 1A. Briefly, the day before experimental procedures began, rats were handled by the experimenter and presented with 25 banana pellets in their homecage to reduce neophobia. The following day, rats began pretraining in which they were placed in the conditioning chambers with 3 pellets already in the food cup, and after 5 minutes the house light was turned on and 25 food pellets were delivered one at a time into the magazine on a variable interval 90 s (30-150 s inter-trial interval) schedule for a session lasting approximately 45 minutes. Rats received one pretraining session a day until a majority of rats consumed all pellets in the pretraining sessions, which took 2-3 days. Pavlovian conditioning began the day after pretraining concluded. Each session consisted of 25 pairings of lever insertion for 8s, after which the lever retracted and a food pellet was delivered in the adjacent food cup. The lever was presented on a variable interval 90 s (30-150 s inter-trial interval) schedule, and each session lasted approximately 40 min. Rats underwent one session a day and were trained for either 7 (standard training) or 14 days (extended training) prior to experimental manipulations. All training and testing took place between 08:00 h and 13:00 h.

**Figure 1.**
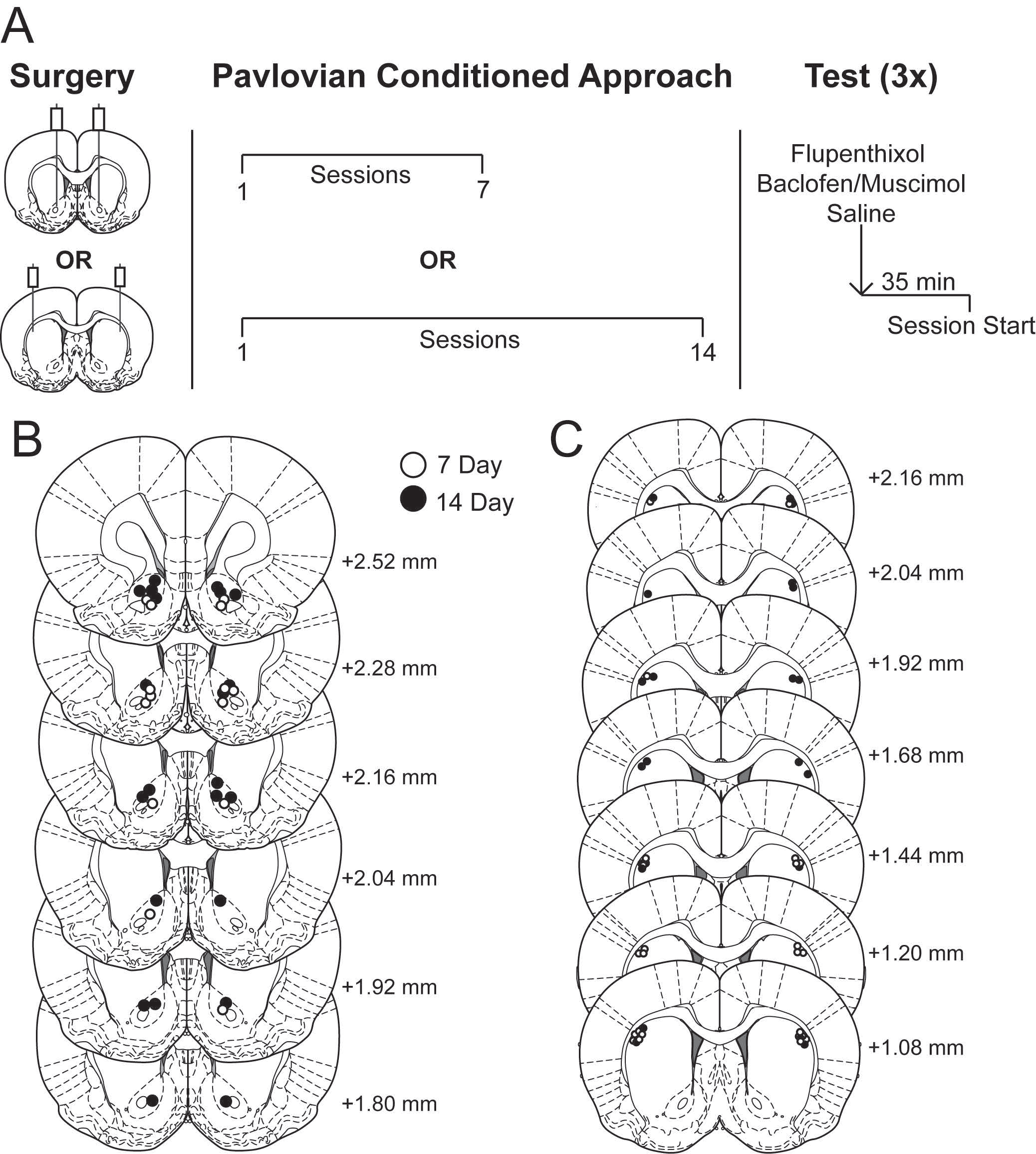
Experimental design and histological verification of microinjector tips. A) Rats received cannulae targeting either the DLS or NAc and were subsequently trained in a Pavlovian conditioning paradigm for 7 or 14 sessions. Each rat received an infusion of saline, the GABA agonists baclofen and muscimol, and the dopamine antagonist flupenthixol across three separate days 35 minutes prior to session start. B) Placements of infuser tips within the nucleus accumbens core. C) Placements of infuser tips within the dorsal lateral striatum. Brain images adapted from Paxinos and Watson (2007) and numbers indicate distance from bregma in millimeters.

Sign-and goal-tracking behavior was quantified using a Pavlovian conditioned approach index score (Meyer et al., 2012). The index score takes into account a rat’s preference to engage with the lever versus the food cup during a Pavlovian conditioning session by averaging the ratio of lever contacts and food cup entries, the difference in probability of contacting the lever or food cup, and the difference in latency to approach the lever or food cup following lever presentation. This results in an index score between −1.0, representing a “perfect” goal-tracker (GT), and 1.0, a “perfect” sign-tracker (ST). Rats were classified using the averaged index score from the last two training sessions before microinfusions (i.e. sessions 6 and 7 for standard training or sessions 13 and 14 for extended training). Rats with index scores between −1.0 and −0.3 were classified as GTs, −0.29 and 0.29 as intermediate responders (INs) who vacillate between food cup and lever contacts on a given trial, and those with scores from 0.3 to 1.0 as STs.

### 2.5 Infusions

Following the completion of the seventh or fourteenth Pavlovian conditioning session rats were assigned to one of three possible drug infusion orders at random. Rats received the dopamine antagonist flupenthixol (100 mM; Sigma, St Louis, MO), a mixture of the GABA_B_ receptor agonist baclofen and GABA_A_ receptor agonist muscimol (B/M; 0.5/0.05 mM; Sigma, St Louis, MO), or saline vehicle prior to the eighth, tenth, or twelfth session (standard training) or the fifteenth, seventeenth, or nineteenth session (extended training). Rats were trained in the sessions between infusions but no infusions or manipulations were made. In order to familiarize rats with the infusion procedure and to check cannula patency rats were brought to the procedure room where infusions would occur following their last Pavlovian training session, they were held by the experimenter, stylets were removed, an infuser was briefly inserted into each cannula and removed. Stylets were then replaced and rats were returned to their home cage. Infusions into the DLS were made in a volume of 0.3 μL over 60 s via 33 gauge infusers and into the NAcC in a volume of 0.5 μL over 90 s via 28 gauge infusers via PE50 tubing connected to 10 μL Hamilton syringes secured in a Harvard Instruments motorized pump. Infusers extended 3 mm past the guide cannula for the DLS (final DV −4.0 mm) and 2 mm past the guide cannula for the NAcC (final DV −7.0 mm). Rats were held gently during the course of infusions. Infusion lines were marked and monitored to ensure that rats received full delivery of solution. Infusers were removed and wire stylets replaced one minute after the completion of the infusion. Rats were tested for the expression of Pavlovian conditioned approach 35 min following drug infusion (Saunders and Robinson, 2012).

### 2.6 Histology

After the completion of the experiment rats were deeply anesthetized with sodium pentobarbital (Euthasol; 0.5 mL intraperitoneally) and decapitated. Brains were extracted and fixed in 4% paraformaldehyde in 0.1 M NaPB overnight then cryoprotected in 25% sucrose in 0.1 M NaPB. Brains were coronally sectioned on a freezing cryostat at −20 C at a thickness of 50 μm, mounted on Fisher SuperFrost Plus slides, and stained with Cresyl violet (FD Neurotechnologies; Ellicott City, MD). Microinjection sites were verified by mapping their locations onto a rat brain atlas (Paxinos and Watson, 2007).

### 2.7 Statistical Analysis

Data were visualized and analyzed in GraphPad Prism 7 (La Jolla, CA). Repeated measures one-way ANOVAs were conducted to analyze the impacts of treatment on lever-and food cup-directed behaviors with α=0.05 for all analyses. Effect sizes were calculated as generalized eta squareds (ηG^2^) using the procedures in Olejnik and Algina (2003). When significant effects were detected post hoc comparisons were performed with Dunnet’s procedure to compare each treatment to saline. In some cases the effect of treatment had a critical non-significant effect, and we followed these analyses by calculating a Bayes factor using the freely available JASP software (v0.8.0.1; https://jasp-stats.org/) to show support for the null hypotheses (Gallistel, 2009). Based on the findings of Saunders and Robinson (Saunders and Robinson, 2012) we had a priori planned comparisons for the effects of flupenthixol versus saline in the nucleus accumbens core on sign-tracking.

## 3. Results

### 3.1 Histological Verification of Cannulae Placements

Three rats were excluded for cannulae failing to be patent and seven rats were excluded for illness or headcap loss prior to testing. The final number of rats for each group was DLS standard training (n=17; n=11 ST; n=3 GT; n=3 IN), DLS extended training (n=15; n=13 ST; n=2 GT), NAcC standard training (n=13; n=7 ST; n=3 IN; n=3 GT), and NAcC extended training (n=15; n=11 ST; n=2 IN; n=2 GT). Due to the low number of GT and IN rats across the four groups, only data from rats meeting the criterion to be classified as ST were analyzed. The locations of infuser tips for rats included in this study are shown in Figure 1B for NAcC and Figure 1C for DLS.

### 3.3 Nucleus Accumbens Core Contributions to Sign-tracking Following Standard Training

In agreement with the findings of Saunders and Robinson (2012), infusion of the dopamine antagonist, flupenthixol, reduced the vigor of lever pressing (F_2,6_=15.96; p<0.01; ηG^2^=0.19; Dunnet’s test p<0.01; Fig. 2A) and increased the latency to approach the lever following presentation (F_2,6_=4.504; p=0.05; ηG^2^=0.09; Dunnet’s test p=0.02; Fig 2C). There was a trend for the treatment reducing probability of lever approach (F_2,6_=4.133; p=0.07; ηG^2^=0.25; Dunnet’s test p=0.07; Fig. 2B). Reversible inactivation with baclofen and muscimol was without effect on sign-tracking (Dunnet’s test all p>0.22). In contrast, reversible inactivation increased the total number of non-specific food cup entries made during the inter-trial interval (F_2,6_=8.261; p=0.03; ηG^2^=0.43; Dunnet’s test p=0.03; Fig. 2D), whereas flupenthixol did not (Dunnet’s test p=0.99).

**Figure 2.**
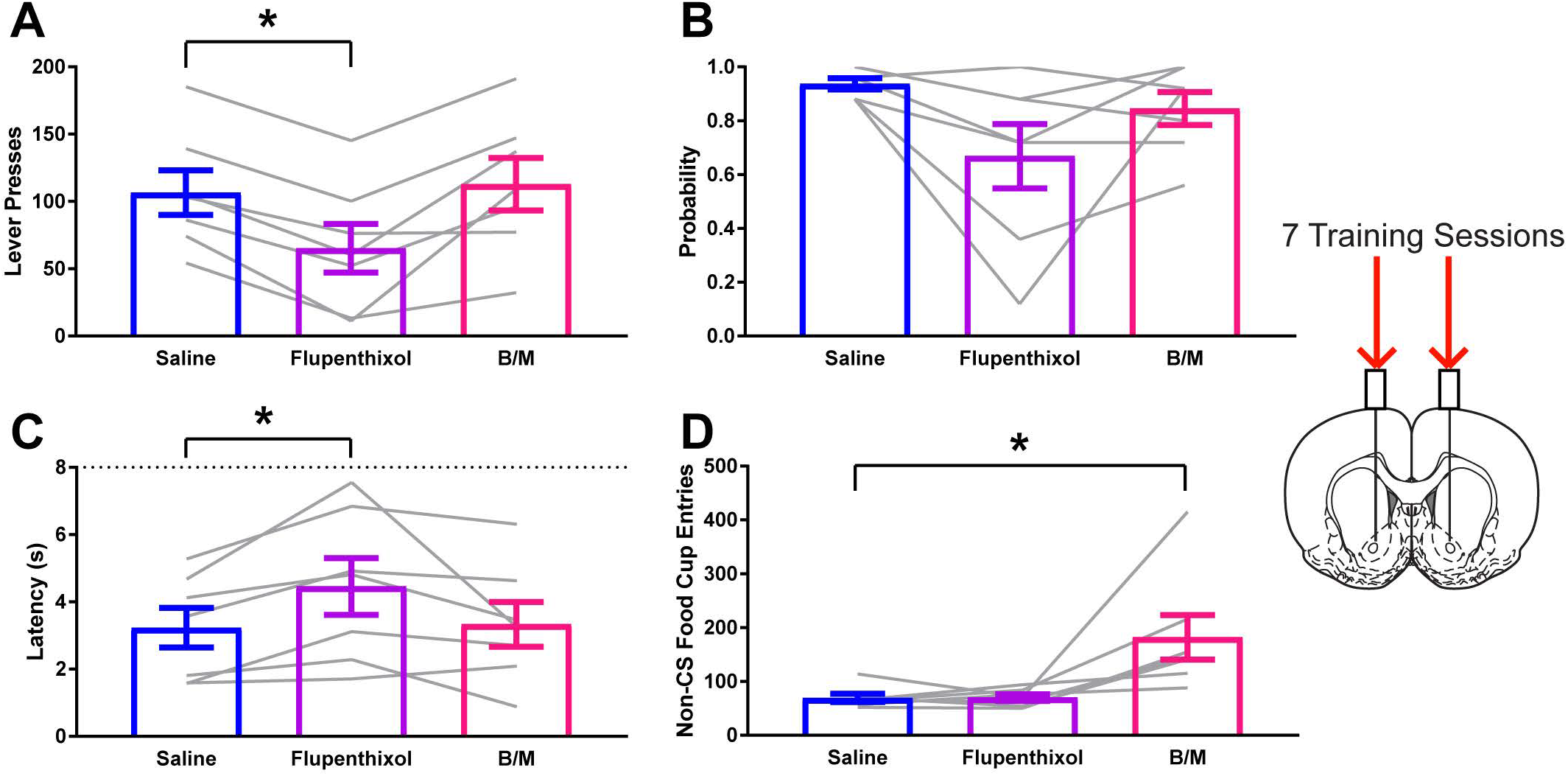
Impact of dopamine antagonism or reversible inactivation of the nucleus accumbens core on sign-tracking behavior following standard training. A) Lever press behavior at test for STs. B) Probability to approach the lever at test for STs. C) Latency to approach the lever following presentation at test for STs. D) Entries into the food cup in the inter-trial interval at test for STs. Data are presented as mean ± S.E.M with an overlay of individual subjects. Brain image adapted from Paxinos and Watson (2007). * p<0.05 for Dunnet’s test comparison.

### 3.3 Nucleus Accumbens Core Contributions to Sign-tracking Following Extended Training

Following extended training, the impact of flupenthixol was similar to that after standard training procedures. Flupenthixol reduced the vigor of lever pressing (F_2,10_=7.945; p<0.01; ηG^2^=0.15; Dunnet’s test p<0.01; Fig 3A), decreased probability of contacting the lever (F_2,10_=3.805; p=0.05; ηG^2^=0.04; Dunnet’s test p=0.05; Fig 3B), and increased the latency to contact the lever (F_2,10_=5.224; p=0.03; ηG^2^=0.10; Dunnet’s test p<0.01; Fig 3C). Reversible inactivation was without effect on these sign-tracking measures (Dunnet’s test all p>0.69). In contrast, reversible inactivation produced a robust increase in food cup entries during the inter-trial intervals (F_2,10_=9.973; p<0.01; ηG^2^=0.34; Dunnet’s test p=0.02; Fig 3D), whereas flupenthixol did not impact this behavior (Dunnet’s test p=0.34).

**Figure 3.**
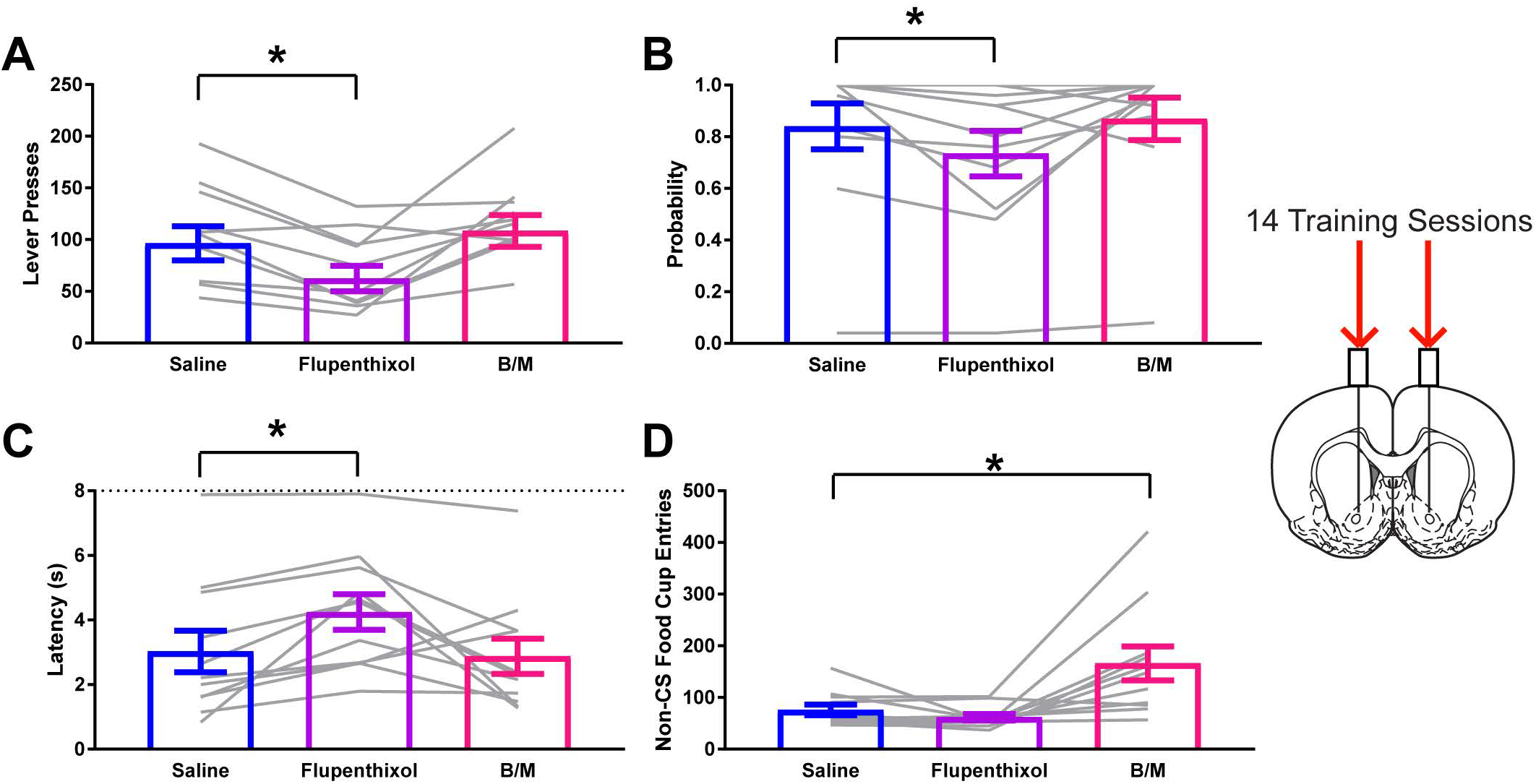
Impact of dopamine antagonism or reversible inactivation of the nucleus accumbens core on sign-tracking behavior after extended training. A) Lever press behavior at test for STs. B) Probability to approach the lever at test for STs. C) Latency to approach the lever following presentation at test for STs. D) Entries into the food cup in the inter-trial interval at test for STs. Data are presented as mean ± S.E.M with an overlay of individual subjects. Brain image adapted from Paxinos and Watson (2007). * p<0.05 for Dunnet’s test comparison.

### 3.4 Dorsal Lateral Striatum Contributions to Sign-tracking Following Standard Training

Neither reversible inactivation of the DLS with the GABA agonists baclofen and muscimol nor dopamine antagonism with flupenthixol impacted the vigor of sign-tracking (Fig. 4A), the probability to approach the lever-cue on a given trial (Fig. 4B), nor the latency to approach the lever-cue following its presentation for STs (Fig. 4C; all F_2,11_<1.49; all ηG^2^<0.02; all p>0.25). Bayesian analyses supported the lack of effect of treatment on sign-tracking as the null hypotheses were 4.44, 3.48, and 2.06 times more likely than the alternative for lever presses, probability, and latency respectively. Neither treatment affected non-specific behavior in the conditioning chamber as measured by food cup entries in the periods outside of lever-cue presentation (Fig. 4D; F_2,11_=1.909; ηG^2^=0.06; p=0.19).

**Figure 4.**
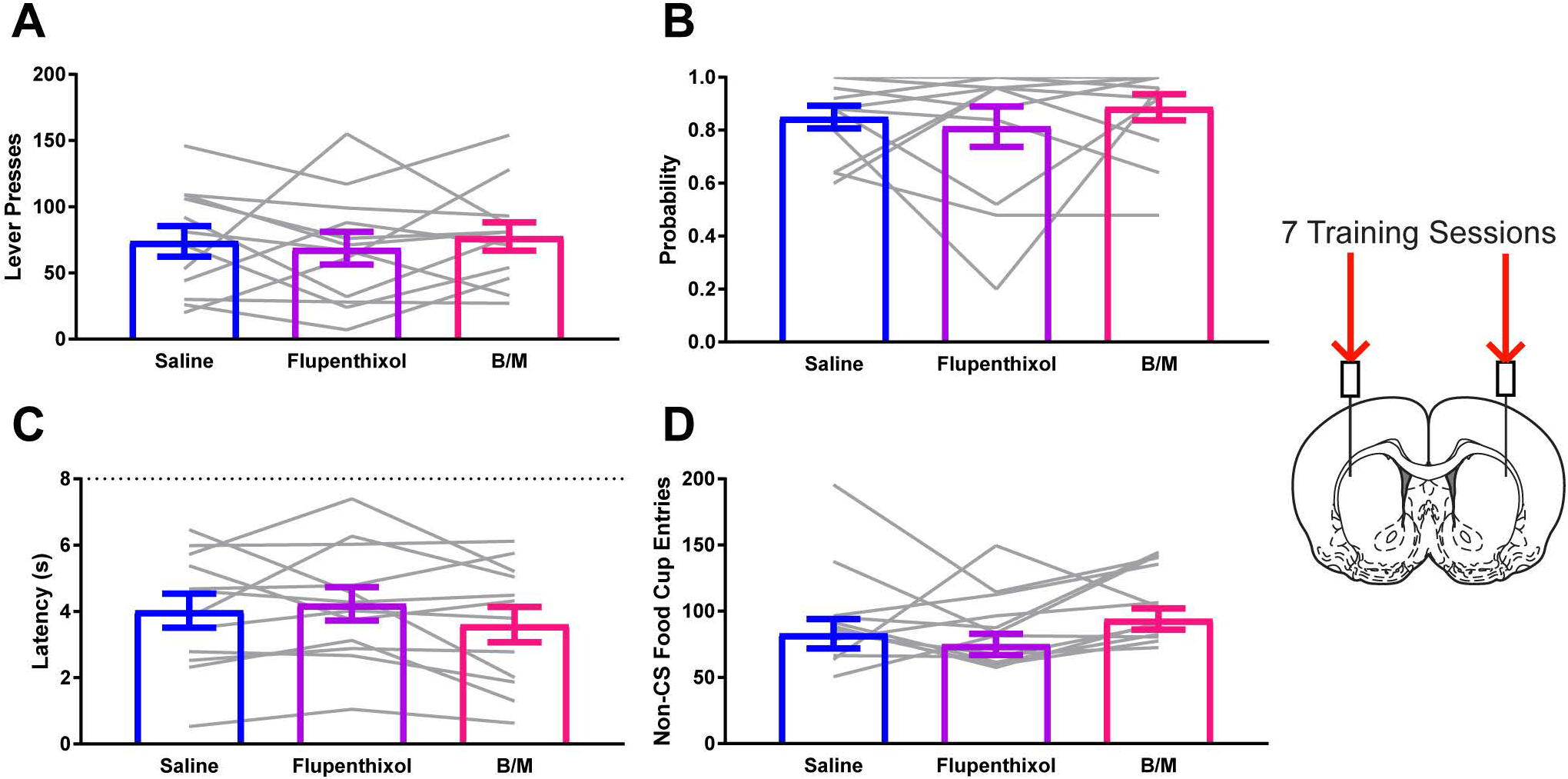
Neither activity in nor dopamine within the dorsal lateral striatum affects sign-tracking following standard training. A) Lever press behavior at test for STs. B) Probability to approach the lever at test for STs. C) Latency to approach the lever following presentation at test for STs. D) Entries into the food cup in the inter-trial interval at test for STs. Data are presented as mean ± S.E.M with an overlay of individual subjects. Brain image adapted from Paxinos and Watson (2007).

### 3.5 Dorsal Lateral Striatum Contributions to Sign-tracking Following Extended Training

Neither reversible inactivation nor dopamine antagonism within the DLS had a significant impact on sign-tracking behavior for STs (Fig 5A-C; all F_2,10_<2.503; all ηG^2^<0.05; all p>0.14). In support of this non-significant effect Bayesian analysis indicated that the null hypothesis was 2.11, 1.1, and 1.2 times more likely than the alternative for lever presses, probability, and latency respectively. There was not a significant impact of either manipulation on non-specific food cup entries (Fig. 5D; F_2,10_=1.444; ηG^2^=0.02; p=0.26).

**Figure 5.**
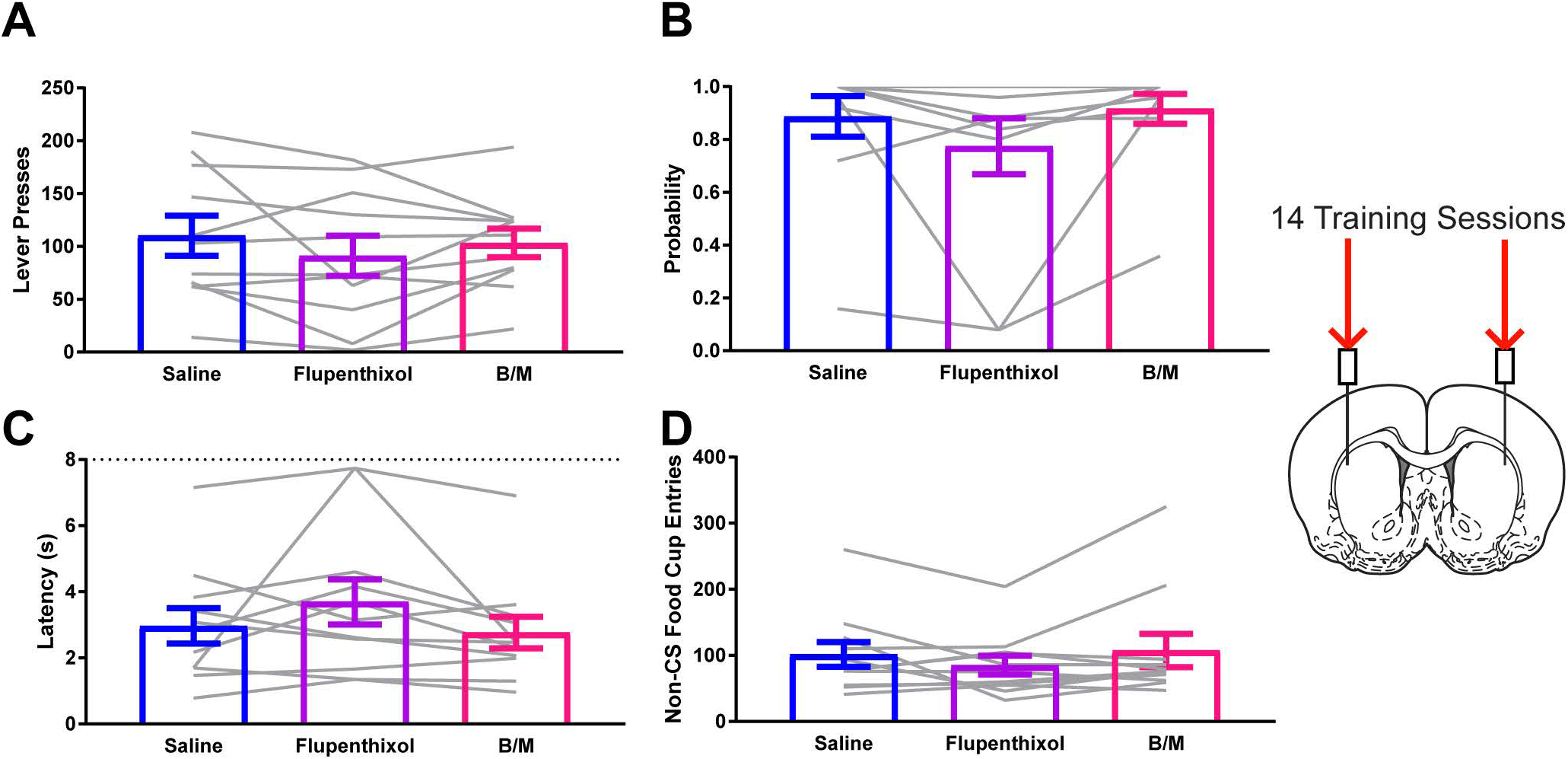
Neither activity in nor dopamine within the dorsal lateral striatum affects sign-tracking after extended training. A) Lever press behavior at test for STs. B) Probability to approach the lever at test for STs. C) Latency to approach the lever following presentation at test for STs. D) Entries into the food cup in the inter-trial interval at test for STs. Data are presented as mean ± S.E.M with an overlay of individual subjects. Brain image adapted from Paxinos and Watson (2007).

## 4. Discussion

We assessed the contributions of both general neural activity and dopamine signaling within the DLS and NAcC to the expression of a sign-tracking conditioned response. Results indicate that, regardless of extended training, functional dopamine signaling in the NAcC, but not DLS, remains critical to the proper execution of a sign-tracking conditioned response. Additionally, we also show that reversible inactivation of either the NAcC or the DLS has minimal impact on sign-tracking behavior at either time point. These data suggest that dopamine’s function in the NAcC remains critical for sign-tracking and there is not a switch in the striatal systems regulating Pavlovian conditioned cue approach with extended training.

Sign-tracking, or Pavlovian conditioned cue approach, is dependent on the attribution of incentive salience to reward-paired cues (Robinson and Flagel, 2009). This is in contrast to goal-tracking, or Pavlovian conditioned goal approach, which does not reflect the attribution of incentive salience to reward-paired cues (Robinson and Flagel, 2009). The attribution of incentive salience is a dopamine dependent process, as systemic blockade of dopamine signaling prevents the acquisition of a sign-but not a goal-tracking conditioned response (Flagel et al., 2011b; Chow et al., 2016; Scülfort et al., 2016). Further, dopamine in the nucleus accumbens core tracks the attribution of incentive salience and dopamine antagonism in the NAcC impairs a sign-tracking, but not goal-tracking, conditioned response (Flagel et al., 2011b; Saunders and Robinson, 2012; Clark et al., 2013). Clark et al., (2013) trained sign-tracking rats while monitoring dopamine efflux in the NAcC for many sessions. Cue-evoked dopamine release, representative of the attribution of incentive salience, peaked as rats reached asymptotic performance, but, past this point with extended training, the signal decayed to levels similar to the initial session before learning occurred. However, there was no direct functional assessment of the decrease Clark et al., (2013) observed in the NAcC, so it has remained unclear if there are behavioral implications to the diminished signal. Although we cannot be certain that our rats indeed showed similar decreases in cue-evoked dopamine release following extended training, we used near-identical procedures as those in Clark et al., (2013), and found dopamine antagonism in the NAcC after extended training reduced the degree to which rats sign-tracked. These data replicate and extended the initial findings of Saunders and Robinson (2012), and suggest dopamine signaling in the NAcC is critical for sign-tracking across a diversity of timescales.

Dopaminergic projections to the striatum exhibit a spiraling network. Projections move ventral and medial to dorsal and lateral in the midbrain, and these terminate in ventral medial to dorsal lateral striatum respectively (Haber et al., 2000). Following extended cocaine self-administration, non-contingent presentation of the cue associated with cocaine delivery elicits dopamine release in the DLS in accompaniment with a decrease in the NAcC (Willuhn et al., 2012, 2014). This switch has been suggested to be critical for escalation of cocaine intake and the transition to addiction, and indeed blockade of dopamine in the DLS is able to dampen cocaine self-administration (Belin and Everitt, 2008; Everitt and Robbins, 2013). These findings suggest that with extended experience the incentive salience attributed to drug-associated cues can recruit systems outside the NAcC. Pharmacologically increasing dopaminergic tone within the DLS has been shown to increase sign-tracking in sign-tracker rats and goal-tracking in goal-tracker rats suggesting increasing dopamine generally increases the vigor of conditioned approach (DiFeliceantonio and Berridge, 2016). However, here we show that blockade of dopamine signaling within the same regions of the DLS that were targeted in the aforementioned studies is without effect on sign-tracking following standard or extended training. Thus, increasing dopamine signaling in the DLS is sufficient to enhance the vigor of Pavlovian conditioned approach behavior but does not appear necessary for its expression. Although it is possible our extended training regimen of 14 sessions prior to manipulations was not sufficient to recruit DLS systems, we favor an interpretation in support of functional distinctions between ventral and dorsal striatal loops, namely that recruitment of dopamine signaling in the dorsal striatum requires the direct linking of an animal’s actions to an outcome through instrumental conditioning. This could serve to explain the dopaminergic response to non-contingent cocaine-associated cues in the DLS as those cues signaled with the completion of a cocaine-seeking instrumental response and subsequent cocaine delivery.

As instrumental actions are performed consistently over time, a switch occurs in which responding is no longer affected based on the value of the reward or goal the action would procure, producing behavior that is habitual and based only on the presentation of the reward-associated stimuli (Yin and Knowlton, 2006). Antagonism of dopamine and glutamate or agonism of GABA signaling within the DLS abolishes the habit-like stimulus-response features of instrumental responding and restores sensitivity to manipulations of outcome value (Yin et al., 2006; Corbit et al., 2012, 2014). In contrast, manipulations made in the dorsal medial striatum produce behaviors that are habitual, as assessed by insensitivity to change in outcome value (Yin et al., 2005; Corbit et al., 2012). This has led to the suggestion that medial and lateral components of the dorsal striatum are in a push-pull dynamic between goal-directed and habitual processes (Yin and Knowlton, 2006; Balleine et al., 2009). Interestingly, sign-tracking, a Pavlovian conditioned response, shares many features of instrumental behaviors that have been described as habitual. For example, sign-tracking is resistant to outcome devaluation (Morrison et al., 2015; Nasser et al., 2015; Patitucci et al., 2016) and extinction (Ahrens et al., 2016), it persists following response-dependent omission of reward (Chang and Smith, 2016), and behavioral inflexibility predicts the degree to which a rat sign-tracks (Nasser et al., 2015). In addition, sign-trackers show greater levels of c-fos protein and mRNA in the dorsal striatum following presentations of a food-cue compared to goal-trackers, suggesting the attribution of incentive salience engages dorsal striatal circuits (Flagel et al., 2011a; Yager et al., 2015).

Activity within the dorsal striatum has previously been shown to contribute minimally to Pavlovian conditioned approach behavior. Corbit and Janak (2007) demonstrated that inactivation of either the medial or lateral portions of the dorsal striatum left Pavlovian conditioned approach driven by a non-localizable auditory cue intact. Blockade of dopamine or reversible inactivation within the dorsal medial striatum is also without effect on sign-or goal-tracking behaviors (O’Donnell, 2014). To our surprise, inactivation of the DLS in the current study was without effect on sign-tracking. This suggests that neither activity nor dopamine signaling in the two main subregions of the dorsal striatum are necessary for Pavlovian conditioned approach, be it to a localizable cue or the location of reward delivery. However, it was recently shown that lesions of the DLS prevented the acquisition of a sign-tracking response, yet this result may be attributable to a motor impairment preventing engagement with the lever (Naeem and White, 2016). A possibility is that rats with DLS lesions in the study by Naeem and White (2016) approached the lever but did not contact it, as has been shown to be the case for sign-tracking following extinction (Chang and Smith, 2016) or to drug-paired Pavlovian cues (Flagel et al., 2010; Yager and Robinson, 2013, 2015; Yager et al., 2015). Taken together, it does not appear that the dorsal striatum is critical for the expression of Pavlovian conditioned approach irrespective of the attribution of incentive salience to a Pavlovian reward-predictive cue.

In contrast to the DLS, inactivation of the NAcC impairs goal-tracking to an auditory cue (Blaiss and Janak, 2009). Contrary to our expectations, inactivation of the NAcC did not prevent the expression of a sign-tracking conditioned response, despite functional dopamine signaling in this region being critical for this behavior. These data, however, are in line with the finding that lesions of either the nucleus accumbens core or shell do not affect sign-tracking to a lever (Chang and Holland, 2013 but see Chang et al., 2012). In contrast to the NAcC, inactivation of the nucleus accumbens shell with baclofen and muscimol disinhibits behavior as evidenced by increasing non-specific behaviors towards available manipulanda in the conditioning chamber (Blaiss and Janak, 2009; Millan et al., 2015). Baclofen and muscimol within the NAcC resulted in an increase in non-specific behavior as rats greatly increased their entries into the food cup during inter-trial intervals. Although our infusions were targeted to the nucleus accumbens core, and only rats with tips within this region were included, a possibility is that the drugs spread into the shell during the incubation period. However, AMPA-receptor antagonism in the nucleus accumbens core increases sign-tracking to a lever never paired with reward without affecting sign-tracking to a reward-paired lever suggesting behavioral disinhibition can be produced by altering excitation-inhibition dynamics within the NAcC (Di Ciano et al., 2001). Here we used only one lever to replicate and compare our findings to previous studies, but perhaps if we had adopted the use of a control lever we would have observed the same effect with GABA agonism. Nonetheless, reversible inactivation of the NAcC with baclofen and muscimol did not directly impair sign-tracking but decreased discriminatory responding in the conditioning chamber.

Identification of sign-tracking as a behavioral phenotype has provided critical insights into the neurobiology of incentive salience attribution and has important implications for substance use disorders (Robinson et al., 2014). Sign-tracking shares features of instrumental habits such as resistance to outcome devaluation and resistance to extinction, suggesting a possible overlap in the systems mediating instrumental habits and sign-tracking. However, alterations of dopamine signaling or activity within the DLS, the region implicated in habit-like responding, were without effect on sign-tracking. Together these data are consistent with a view that the NAcC remains a critical neural center for the attribution of incentive salience even extended experience and for related processes like incentive sensitization (Robinson and Berridge, 1993; Berridge and Robinson, 1998; Berridge, 2007). These findings also suggest that recruitment of dorsal striatal systems depends on instrumental processes linking an organism’s behavior directly to reward procurement as opposed to a Pavlovian setting where reward occurs independent of behavior. It will be important in the future to resolve the inputs, projection target, and cellular phenotype of those neurons in the NAcC responsible for sign-tracking to better understand the neurobiology of incentive salience and guide potential treatments for substance use disorders.

## Acknowledgements

This work was supported by NIH grant R01 DA035943 from the National Institute on Drug Abuse to P.H.J. We thank Alex Haimbaugh for excellent technical assistance. We would also like to thank Drs. Shelly Flagel, Joshua Haight, Peter Holland, Ronald Keiflin, E. Zayra Millan, Jocelyn Richard, Benjamin Saunders, and Youna Vandaele as well as Brittany Kuhn for inputs on study design, discussions of early results, and for their continued support and encouragement.

## Author Contributions

KMF designed and carried out experiments, analyzed data, and wrote the manuscript. PHJ designed experiments and wrote the manuscript.

## Data Accessibility

Data will be made available upon request.

## References

Ahrens AM, Singer BF, Fitzpatrick CJ, Morrow JD, Robinson TE (2016) Rats that sign-track are resistant to Pavlovian but not instrumental extinction. Behav Brain Res 296:418–430.

Anderson RI, Bush PC, Spear LP (2013) Environmental manipulations alter age differences in attribution of incentive salience to reward-paired cues. Behav Brain Res 257:83–89.

Balleine BW, Liljeholm M, Ostlund SB (2009) The integrative function of the basal ganglia in instrumental conditioning. Behav Brain Res 199:43–52.

Belin D, Everitt BJ (2008) Cocaine seeking habits depend upon dopamine-dependent serial connectivity linking the ventral with the dorsal striatum. Neuron 57:432– 441.

Berridge KC (2001) Reward Learning: Reinforcement, Incentives and Expectations. In: Psychology of Learning and Motivation: Advances in Research and theory (Medin DL, ed), pp 223–278. Academic Press.

Berridge KC (2007) The debate over dopamine’s role in reward: the case for incentive salience. Psychopharmacology (Berl) 191:391–431.

Berridge KC (2012) From prediction error to incentive salience: mesolimbic computation of reward motivation. Eur J Neurosci 35:1124–1143.

Berridge KC, Robinson TE (1998) What is the role of dopamine in reward: hedonic impact, reward learning, or incentive salience? Brain Res Brain Res Rev 28:309– 369.

Blaiss CA, Janak PH (2009) The nucleus accumbens core and shell are critical for the expression, but not the consolidation, of Pavlovian conditioned approach. Behav Brain Res 200:22–32.

Boakes R (1977) Performance on learning to associate a stimulus with positive reinforcement. In: Operant-Pavlovian interactions (Davis H, Hurwitz H, eds), pp 67–97. Hillsdale, NJ: Lawrence Erlbaum Associates.

Cardinal RN, Parkinson JA, Hall J, Everitt BJ (2002) Emotion and motivation: the role of the amygdala, ventral striatum, and prefrontal cortex. Neurosci Biobehav Rev 26:321–352.

Chang SE, Holland PC (2013) Effects of nucleus accumbens core and shell lesions on autoshaped lever-pressing. Behav Brain Res 256:36–42.

Chang SE, Smith KS (2016) An omission procedure reorganizes the microstructure of sign-tracking while preserving incentive salience. Learn Mem 23:151–155.

Chang SE, Wheeler DS, Holland PC (2012) Roles of nucleus accumbens and basolateral amygdala in autoshaped lever pressing. Neurobiol Learn Mem 97:441–451.

Chow JJ, Nickell JR, Darna M, Beckmann JS (2016) Toward isolating the role of dopamine in the acquisition of incentive salience attribution. Neuropharmacology 109:320–331.

Clark JJ, Collins AL, Sanford CA, Phillips PEM (2013) Dopamine encoding of Pavlovian incentive stimuli diminishes with extended training. J Neurosci 33:3526–3532.

Corbit LH, Janak PH (2007) Inactivation of the lateral but not medial dorsal striatum eliminates the excitatory impact of Pavlovian stimuli on instrumental responding. J Neurosci 27:13977–13981.

Corbit LH, Nie H, Janak PH (2012) Habitual alcohol seeking: time course and the contribution of subregions of the dorsal striatum. Biol Psychiatry 72:389–395.

Corbit LH, Nie H, Janak PH (2014) Habitual responding for alcohol depends upon both AMPA and D2 receptor signaling in the dorsolateral striatum. Front Behav Neurosci 8:301.

Di Ciano P, Cardinal RN, Cowell RA, Little SJ, Everitt BJ (2001) Differential involvement of NMDA, AMPA/kainate, and dopamine receptors in the nucleus accumbens core in the acquisition and performance of pavlovian approach behavior. J Neurosci 21:9471–9477.

DiFeliceantonio AG, Berridge KC (2016) Dorsolateral neostriatum contribution to incentive salience: opioid or dopamine stimulation makes one reward cue more motivationally attractive than another. Eur J Neurosci 43:1203–1218.

Everitt BJ, Robbins TW (2013) From the ventral to the dorsal striatum: devolving views of their roles in drug addiction. Neurosci Biobehav Rev 37:1946–1954.

Flagel SB, Cameron CM, Pickup KN, Watson SJ, Akil H, Robinson TE (2011a) A food predictive cue must be attributed with incentive salience for it to induce c-fos mRNA expression in cortico-striatal-thalamic brain regions. Neuroscience 196:80–96.

Flagel SB, Clark JJ, Robinson TE, Mayo L, Czuj A, Willuhn I, Akers CA, Clinton SM, Phillips PEM, Akil H (2011b) A selective role for dopamine in stimulus-reward learning. Nature 469:53–57.

Flagel SB, Robinson TE (2017) Neurobiological basis of individual variation in stimulus-reward learning. Current Opinion in Behavioral Sciences 13:178–185.

Flagel SB, Robinson TE, Clark JJ, Clinton SM, Watson SJ, Seeman P, Phillips PEM, Akil H (2010) An animal model of genetic vulnerability to behavioral disinhibition and responsiveness to reward-related cues: implications for addiction. Neuropsychopharmacology 35:388–400.

Fraser KM, Haight JL, Gardner EL, Flagel SB (2016) Examining the role of dopamine D2 and D3 receptors in Pavlovian conditioned approach behaviors. Behav Brain Res 305:87–99.

Gallistel CR (2009) The importance of proving the null. Psychol Rev 116:439–453.

Haber SN, Fudge JL, McFarland NR (2000) Striatonigrostriatal pathways in primates form an ascending spiral from the shell to the dorsolateral striatum. J Neurosci 20:2369–2382.

Haight JL, Fraser KM, Akil H, Flagel SB (2015) Lesions of the paraventricular nucleus of the thalamus differentially affect sign-and goal-tracking conditioned responses. Eur J Neurosci 42:2478–2488.

Hearst E, Jenkins H (1974) Sign-tracking?: the stimulus-reinforcer relation and directed action. Austin, TX: Proceedings of the Psychonomic Society.

Hollerman JR, Schultz W (1998) Dopamine neurons report an error in the temporal prediction of reward during learning. Nat Neurosci 1:304–309.

Keiflin R, Janak PH (2015) Dopamine prediction errors in reward learning and addiction: from theory to neural circuitry. Neuron 88:247–263.

Meyer PJ, Lovic V, Saunders BT, Yager LM, Flagel SB, Morrow JD, Robinson TE (2012) Quantifying individual variation in the propensity to attribute incentive salience to reward cues. PLoS ONE 7:e38987.

Millan EZ, Reese RM, Grossman CD, Chaudhri N, Janak PH (2015) Nucleus Accumbens and Posterior Amygdala Mediate Cue-Triggered Alcohol Seeking and Suppress Behavior During the Omission of Alcohol-Predictive Cues. Neuropsychopharmacology 40:2555–2565.

Morrison SE, Bamkole MA, Nicola SM (2015) Sign Tracking, but Not Goal Tracking, is Resistant to Outcome Devaluation. Front Neurosci 9:468.

Naeem M, White NM (2016) Parallel learning in an autoshaping paradigm. Behav Neurosci 130:376–392.

Nasser HM, Chen Y-W, Fiscella K, Calu DJ (2015) Individual variability in behavioral flexibility predicts sign-tracking tendency. Front Behav Neurosci 9:289.

O’Donnell E (2014) Dorsomedial Striatal Control of Cue-Directed Versus Goal-Directed Pavlovian Approach Behavior. Available at: https://deepblue.lib.umich.edu/handle/2027.42/102760.

Olejnik S, Algina J (2003) Generalized eta and omega squared statistics: measures of effect size for some common research designs. Psychol Methods 8:434–447.

Patitucci E, Nelson AJD, Dwyer DM, Honey RC (2016) The origins of individual differences in how learning is expressed in rats: A general-process perspective. Journal of experimental psychology Animal learning and cognition 42:313–324.

Paxinos G, Watson C (2007) The Rat Brain in Stereotaxic Coordinates, 6th ed. New York: Academic Press.

Robinson TE, Berridge KC (1993) The neural basis of drug craving: an incentive-sensitization theory of addiction. Brain Res Brain Res Rev 18:247–291.

Robinson TE, Flagel SB (2009) Dissociating the predictive and incentive motivational properties of reward-related cues through the study of individual differences. Biol Psychiatry 65:869–873.

Robinson TE, Yager LM, Cogan ES, Saunders BT (2014) On the motivational properties of reward cues: Individual differences. Neuropharmacology 76 Pt B:450–459.

Saunders BT, Robinson TE (2010) A cocaine cue acts as an incentive stimulus in some but not others: implications for addiction. Biol Psychiatry 67:730–736.

Saunders BT, Robinson TE (2012) The role of dopamine in the accumbens core in the expression of Pavlovian-conditioned responses. Eur J Neurosci 36:2521–2532.

Saunders BT, Robinson TE (2013) Individual variation in resisting temptation: implications for addiction. Neurosci Biobehav Rev 37:1955–1975.

Saunders BT, Yager LM, Robinson TE (2013) Cue-evoked cocaine “craving”: role of dopamine in the accumbens core. J Neurosci 33:13989–14000.

Schultz W, Dayan P, Montague PR (1997) A neural substrate of prediction and reward. Science 275:1593–1599.

Scülfort SA, Bartsch D, Enkel T (2016) Dopamine antagonism does not impair learning of Pavlovian conditioned approach to manipulable or non-manipulable cues but biases responding towards goal tracking. Behav Brain Res 314:1–5.

Steinberg EE, Keiflin R, Boivin JR, Witten IB, Deisseroth K, Janak PH (2013) A causal link between prediction errors, dopamine neurons and learning. Nat Neurosci 16:966–973.

Vollstädt-Klein S, Wichert S, Rabinstein J, Bühler M, Klein O, Ende G, Hermann D, Mann K (2010) Initial, habitual and compulsive alcohol use is characterized by a shift of cue processing from ventral to dorsal striatum. Addiction 105:1741–1749.

Willuhn I, Burgeno LM, Everitt BJ, Phillips PEM (2012) Hierarchical recruitment of phasic dopamine signaling in the striatum during the progression of cocaine use. Proc Natl Acad Sci U S A 109:20703–20708.

Willuhn I, Burgeno LM, Groblewski PA, Phillips PEM (2014) Excessive cocaine use results from decreased phasic dopamine signaling in the striatum. Nat Neurosci 17:704–709.

Yager LM, Pitchers KK, Flagel SB, Robinson TE (2015) Individual variation in the motivational and neurobiological effects of an opioid cue. Neuropsychopharmacology 40:1269–1277.

Yager LM, Robinson TE (2013) A classically conditioned cocaine cue acquires greater control over motivated behavior in rats prone to attribute incentive salience to a food cue. Psychopharmacology (Berl) 226:217–228.

Yager LM, Robinson TE (2015) Individual variation in the motivational properties of a nicotine cue: sign-trackers vs. goal-trackers. Psychopharmacology (Berl) 232:3149–3160.

Yin HH, Knowlton BJ (2006) The role of the basal ganglia in habit formation. Nat Rev Neurosci 7:464–476.

Yin HH, Knowlton BJ, Balleine BW (2006) Inactivation of dorsolateral striatum enhances sensitivity to changes in the action-outcome contingency in instrumental conditioning. Behav Brain Res 166:189–196.

Yin HH, Ostlund SB, Knowlton BJ, Balleine BW (2005) The role of the dorsomedial striatum in instrumental conditioning. Eur J Neurosci 22:513–523.

